# Binding of the TRF2 iDDR motif to RAD50 highlights a convergent evolutionary strategy to inactivate MRN at telomeres

**DOI:** 10.1101/2023.03.25.534200

**Authors:** Freddy Khayat, Majedh Alshmery, Mohinder Pal, Antony W. Oliver, Alessandro Bianchi

## Abstract

Telomeres protect chromosome ends from unscheduled DNA repair, including from the MRN (MRE11, RAD50, NBS1) complex, which plays a critical role in the processing of double-stranded DNA breaks (DSBs). MRN orchestrates activation of the ATM kinase in the cellular DNA damage response (DDR), promotes DNA end-tethering aiding the nonhomologous end joining (NHEJ) pathway, and initiates DSB resection through the MRE11 nuclease. A previously identified protein motif (MIN, for MRN inhibitor) downregulates MRN activity via binding to RAD50 and has independently arisen at least twice, through convergent evolution of telomeric proteins Rif2 and Taz1, in budding and fission yeast respectively. We now provide a third example of convergent evolution for this binding mechanism for MRN at telomeres, by demonstrating that the iDDR motif of the human shelterin protein TRF2 binds to human RAD50 at the same site engaged by the MIN motif in the yeast proteins, despite lacking sequence homology. Modelling for the human CtIP interaction with RAD50 (necessary for activation of MRE11), and for the budding and fission yeast counterparts Sae2 and Ctp1, indicates that the interaction is mutually exclusive with binding of the iDDR/MIN motifs, pointing to a conserved mechanism for inhibition of MRN nuclease activity at telomeres.

## INTRODUCTION

Eukaryotes have evolved specialised structures, the telomeres, to protect the ends of their segmented linear genomes. Protection requires maintenance and replication of the telomeres themselves, by the telomerase enzyme, and concomitant inhibition of DNA repair pathways, such as DNA double-strand break (DSB) repair, that would otherwise compromise integrity of the ends (Lustig, 2019; Lazzerini-Denchi and Sfeir, 2016). To execute these functions telomeres possess a number of conserved features, which include tandem arrays of DNA repeats, mostly configured as double-stranded DNA but with a terminal single-stranded 3’ overhang. These repeats are bound by a set of DNA double- and single-stranded telomeric DNA binding factors that assemble a protein complex at the chromosome ends (Lim and Cech, 2021; de Lange, 2018). Despite these general similarities, the exact protein make-up of this complex is highly variable within eukaryotes, suggesting a scenario where gene duplications and divergence within the telomeric repeats - due to mutations in the telomerase RNA template - have led to diversity in the set, arrangement, and number of telomere associated factors (Červenák et al., 2017; Červenák et al., 2021; Myler et al., 2021; Xue et al., 2017; Lue, 2021). This points to a highly malleable complex where the main feature is the presence of a conserved set of functionalities but within a great degree of architectural variation and structural flexibility, a notion that has been confirmed experimentally (Zinder et al., 2022).

The core set of telomeric factors in metazoans is constituted by shelterin, which in humans is made of six protein factors (de Lange, 2018), whereas in budding yeast the telomeric complex takes a markedly different form (Wellinger and Zakian, 2012). In all cases, however, the action of additional factors - whose function is not primarily telomeric - is co-opted at telomeres. A case in point is represented by the MRN complex itself and the two main kinases involved in the DNA damage response (DDR), ATM and ATR. MRN (MRX in budding yeast, comprising of RAD50, MRE11 and NBS1/Xrs2) acts as the main sensor of DSBs (Syed and Tainer, 2018; Hopfner, 2023) and is responsible for their resection via the endonucleolytic action of the MRE11 nuclease, which requires also the ATPase activity of RAD50, and the CtIP/Sae2/Ctp1 co-factor (Cannavo and Cejka, 2014; Anand et al., 2016; Deshpande et al., 2016). In addition, MRN is required for activation of the ATM/Tel1 kinase (Hailemariam et al., 2019b; Roset et al., 2014; Lee et al., 2013). In yeast, the MRX complex is additionally required for DSB repair by NHEJ (Boulton and Jackson, 1998; Reis et al., 2012), an activity that has been attributed to the ability of the long coiled-coil stemming from the globular head domain of RAD50 to interact intermolecularly and to promote DNA end-tethering (Rotheneder et al., 2023; Park et al., 2017). Although telomeres suppress the ability of MRN to promote DNA repair, in budding yeast MRX/Tel1 also plays an important role in activating telomerase (Nugent et al., 1998) whilst in human cells ATM can similarly promote telomerase action (Tong et al., 2015; Lee et al., 2015). Elegant experiments carried out in yeast have shown that MRX/Tel1 is responsible for resection and telomerase recruitment at telomeres replicated by the leading strand, likely due to the presence of a blunt end after DNA replication (Faure et al., 2010; Soudet et al., 2014). Intriguingly, at mammalian telomeres leading telomere resection and overhang-establishment are not executed by MRN but by the Apollo nuclease (Wu et al., 2010). Telomeres therefore participate in a delicate and balancing act serving to regulate MRN activity.

The telomeric protein complex in the budding yeast *Saccharomyces cerevisiae* is assembled from the DNA binding protein Rap1 which, among other things, recruits the telomere factor Rif2, which originated from duplication of the essential Orc4 gene in a subset of yeast clades (Marcand et al., 2008). Rif2 was shown to be part of a Rap1 pathway leading to inhibition of Tel1 in promoting telomere length (Craven and Petes, 1999), with work from the Longhese laboratory providing an early glimpse of the ability of Rif2 to counteract Tel1, by showing that overexpression of Rif2 is sufficient to suppress hyperactivation of Tel1 (Viscardi et al., 2003). Further work has demonstrated that Rif2 acts to ‘mark’ short telomeres for preferential elongation by telomerase (McGee et al., 2010), likely by counteracting increased activation of Tel1 at shortened telomeres (Hector et al., 2007; Sabourin et al., 2007; Bianchi and Shore, 2007).

More recent work has shed light on a possible mechanism for the control of MRN activity at yeast telomeres by Rif2, through identification of a short amino acid motif within the protein (called MIN, for MRN-inhibitor, or BAT, for blocks the addition of telomeres) that is capable of disabling MRN activity in DNA resection, in DNA repair by NHEJ, and in the activation of Tel1 (Khayat et al., 2021; Roisné-Hamelin et al., 2021; Kaizer et al., 2015; Hailemariam et al., 2019a). The MIN motif acts by binding to the RAD50 subunit, to an exposed β-sheet that is also responsible for making contact with Sae2 (Khayat et al., 2021; Roisné-Hamelin et al., 2021; Marsella et al., 2021). Within the MRN complex, RAD50 fulfils a role as a crucial regulatory subunit, responsible for orchestrating an ATP-driven shift from a ‘resting’ ATP-bound conformation to a ‘cutting’ one that is ADP-bound (Lafrance-Vanasse et al., 2015; Deshpande et al., 2014). While the former is proficient for DNA binding and tethering, as well as for Tel1/ATM activation and for NHEJ, the latter confers nuclease activity (Rotheneder et al., 2023; Käshammer et al., 2019). The MIN motif could in principle disable nucleolytic action by displacing Sae2 from MRN, and counter NHEJ and Tel1 activation by interfering with the ‘resting’ conformation, possibly in light of its ability to increase the rate of ATP hydrolysis by RAD50 (Roisné-Hamelin et al., 2021; Hailemariam et al., 2019a; Khayat et al., 2021; Marsella et al., 2021; Cassani et al., 2016).

Although the importance of MRN in processing telomeric ends under physiological conditions might vary in different organisms, it is clear that MRN has a unique role in activating ATM at deprotected telomeres, for example after loss of shelterin subunit TRF2 (Dimitrova and de Lange, 2009; Deng et al., 2009; Attwooll et al., 2009). TRF2 plays a key role in orchestrating multiple aspects of mammalian telomere protection, associating directly with the MRN complex via a highly conserved amino acid motif (called iDDR, for inhibitor of DDR) which is located in an unstructured region nestled between the TRF2 homodimerization (TRFH) and DNA binding motifs, and which suppresses the DDR at deprotected telomeres (Okamoto et al., 2013). Because TRF2 interacts directly with NBS1 through its TRFH domain, in a manner regulated by NBS1 phosphorylation (Rai et al., 2017), and also with ATM (Karlseder et al., 2004), it remains unclear whether TRF2, or any other shelterin subunit, might act to regulate MRN at telomeres using a strategy similar to that employed by the MIN motif of yeast Rif2. Here we address this question and show that the iDDR motif of TRF2 does indeed bind to RAD50 in a manner similar to that of the MIN motif. This finding represents the third example that we document for the convergent evolution of this mechanism that inhibits the action of MRN at telomeres, thus underscoring it functional relevance.

## MATERIAL AND METHODS

### Plasmids

Plasmids were constructed in pGBKT7 or pGADT7, except for the constructs in Figure 1C, which were made in modified vectors for either N- or C-terminal tagging. All mutants were created by PCR with the desired changes embedded in the PCR primers, followed by incorporation of one or two fragments into the appropriate vectors by Gibson Assembly using home-made reaction mixes (Gibson, 2011). All clones were verified by sequencing.

**Figure 1.**
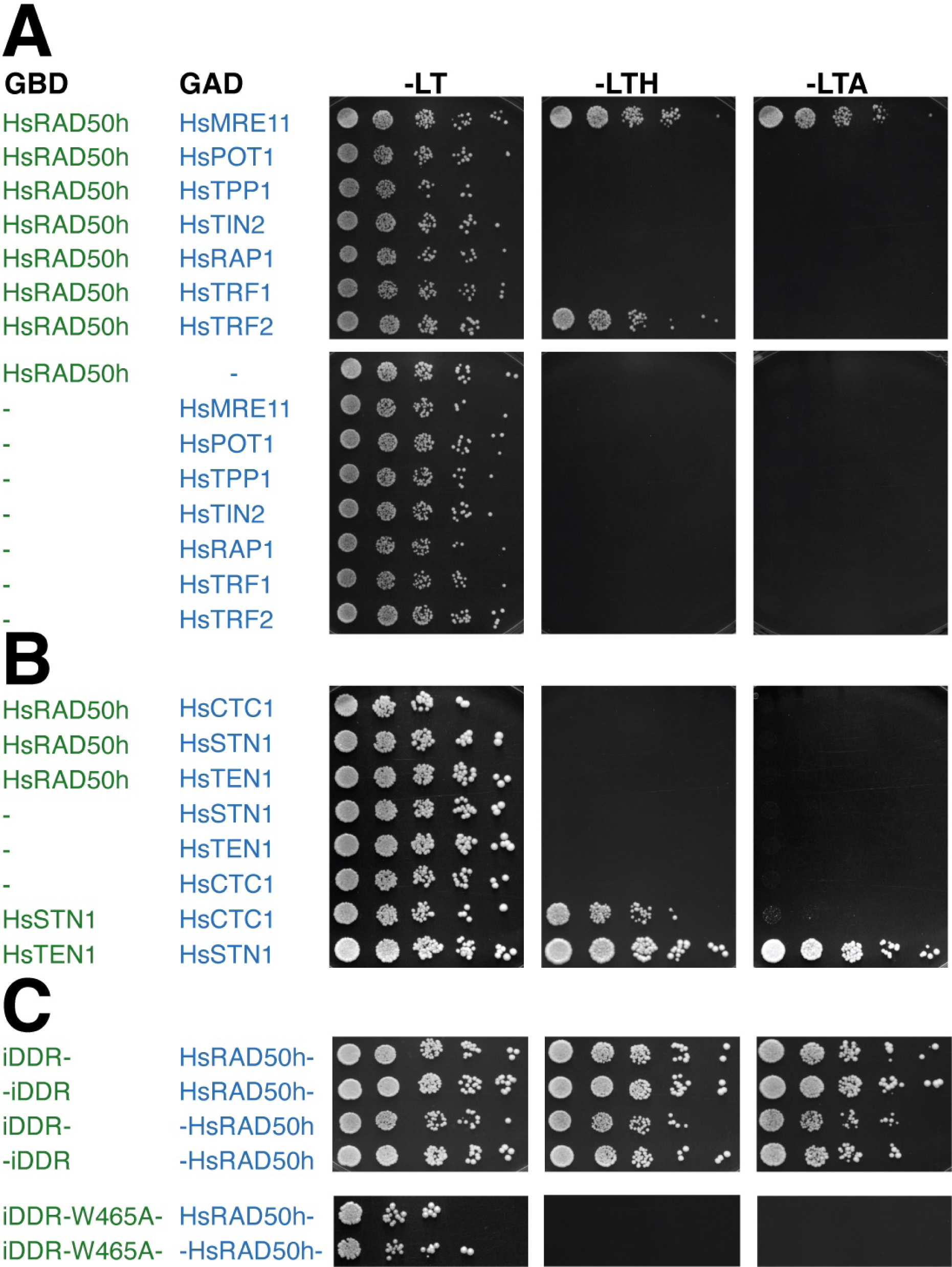
The iDDR motif of TRF2 binds RAD50. (**A**) Yeast two-hybrid analysis of full-length human shelterin subunits fused to the GAD domain of Gal4 for interaction against GBD-HsRAD50 encompassing HsRAD50 amino acids 1-221,1098-1312, separated by an SGSSAGG linker; this HsRAD50 ‘head’ (HsRAD50h) construct lacks the majority of the coiled-coil ‘arm’. Culture dilutions were spotted on plates lacking tryptophan and leucine (-LT) as controls for cell numbers, or tryptophan, leucine and histidine (-LTH) for selection of interaction, or tryptophan, leucine and adenine (-LTA) for more stringent selection for interaction. HsMRE11 was used as a positive control for interaction with HsRAD50h. (**B**) Analysis as in (A) testing the interaction of the CST subunits HsCTC1, HsSTN1 and HsTEN1 with the HsRAD50h construct. (**C**) Analysis as in (A) testing the interaction of the HsTRF2 region spanning amino acids 449-473, which includes iDDR, fused to the GBD of Gal4, either at the N- or C-terminus, with the head domain of human RAD50 (HsRAD50h) fused to either the N- or C-terminu of the Gal4 GAD domain; in all cases, the hyphen at the beginning or the end of the construct names indicates the point of the GBD/GAD fusion. Negative controls (*vs* empty vectors) for both the wild-type TRF2 and RAD50 truncations were performed and did not show growth on selective -LTH and -LTA plates (not shown).

### Yeast two-hybrid analysis

Strains were grown in synthetic complete (SC) minimal medium (2% w/v glucose, 0.67% w/v Yeast Nitrogen Base, 0.2% w/v SC dropout mix, 2% w/v agar, pH 6.0). YNB and SC mixes were from United States Biological. For transformations, the LiAc protocol was followed, with heat shock at 42°C for 45 min. Yeast two hybrid assays were performed by transforming GBD (Gal4 DNA Binding Domain) and GAD (Gal4 Activating Domain) plasmids in the haploid budding yeast strains Y2HGold (YAB2356) and Y187 (YAB2357), respectively. YAB2356: MAT<u>a</u> trp1-901 leu2-3,112 ura3-52 his3-200 gal4Δ gal80Δ LYS2 : : GAL1_UAS_–Gal1_TATA_–His3 GAL2_UAS_–Gal2_TATA_– Ade2 URA3::MEL1_UAS_–Mel1_TATA_ AUR1-C MEL1. YB2357: MATα ura3-52 his3-200 ade2-101 trp1-901 leu2-3,112 gal4Δ gal80Δ met– URA3::GAL1_UAS_–Gal1_TATA_–LacZ MEL1. GBD and GAD transformants were selected on SC medium lacking tryptophan or leucine, respectively. Haploid strains were then mated on rich-medium YPAD plates, and diploids were selected after one day by streaking on plates lacking both tryptophan and leucine. Interactions were assessed by quantification of expression of the HIS3 and ADE2 markers by spotting 5-fold dilutions of liquid cultures on minimal medium lacking, in addition, histidine or adenine, starting with 20,000 cells. Plates were incubated at 30°C for 2-3 (SC-leu-trp) or 3-4 (selective plates) days.

### AlphaFold2 models

All models were generated using ColabFold v1.5.2: AlphaFold2 using MMseqs2 (Mirdita et al., 2022). In each case, 5 amber-relaxed models were generated, with no specified or provided template. The option to use the ‘alphafold2_multimer_v3’ model type was selected. All other parameters were left at their default settings. To reduce computational ‘cost’ the amino acid sequence corresponding to just the head-domain of RAD50 was provided in each case: consisting of the N-terminal portion and a short (∼ 44 amino acid) section of the ascending helix ‘fused’ to a similar section of the descending helix and C-terminal portion by an artificial linking segment consisting of the amino acid sequence ‘GGGGGGSGGGGGGSGGGGGG’.

### Molecular graphics

Figures were generated using either PyMOL (v. 2.5.4) (Schrödinger) (https://pymol.org/2) or ChimeraX (v. 1.1.1) (Pettersen et al., 2021). PyMOL was used to colour models according to the predicted local distance difference test (pLLDT) scores present in the B-factor column of the PDB file generated for each model, using a ‘reverse rainbow spectrum’ ranging from blue (pLDDT score >80, high confidence) to red (pLDDT score < 50, low confidence or intrinsically disordered region). Electrostatic surfaces were calculated using the inbuilt functionality of ChimeraX.

## RESULTS

### The iDDR motif of TRF2 interacts directly with RAD50

We recently reported identification of a short amino acid motif (MIN, for MRN inhibitor) in yeast telomeric proteins that has the ability to bind the RAD50 subunit of the MRN complex and disable the complex’s function. There, we suggested that the MIN motif has evolved at least twice during evolution: in Orc4 in the Saccharomycotina, giving rise to the *Saccharomyces cereviase* (Sc) Rif2 MIN motif and in the *Schizosaccharomyces* genus of the Taphrynomicotina, giving rise to the *Schizosaccharomyces pombe* (Sp) Taz1 MIN motif (Khayat et al., 2021). This identified a potential general vulnerability in MRN and pointed to the possibility that other organisms might have independently evolved the same functionality within their telomeric proteins to regulate the activity of MRN, conceivably though the use of different motifs with unrelated primary sequence. Since we failed to identify the MIN motif in interesting candidates in the human genome by sequence analysis, we decided to investigate the ability of human core telomeric proteins - i.e., members of the shelterin complex - to interact with RAD50, using a yeast two-hybrid approach (Y2H). Our Y2H data reveal that TRF2 is unique among shelterin proteins in its ability to bind RAD50 (Figure 1A), suggesting a direct interaction, given that yeast is a heterologous system for human proteins; this conclusion is in agreement with previous work showing association of TRF2 with MRN and a preferential enrichment of the RAD50 subunit, over MRE11 and NBS1, in TRF2 immunoprecipitates (Okamoto et al., 2013; Zhu et al., 2000). Moreover, no interaction with RAD50 was observed for proteins of the CST (CTC1, STN1, TEN1) complex (Figure 1B). As previous work had attributed the ability of TRF2 to directly bind MRN to the short iDDR motif in the hinge region of TRF2 - at least when inserted in full-length TRF1 (Okamoto et al., 2013) - we checked the ability of iDDR to bind RAD50 on its own. The iDDR motif, spanning TRF2 amino acids (aa) 449-473, was found to be sufficient for interaction with RAD50 in two configurations tested, specifically when it was fused to the GBD motif of Gal4 at either its N- or C-terminus (Figure 1C). Introduction of an Ala mutation at the highly conserved position Trp465 (see Figure 2C below) abolished the interaction (Figure 1C). Taken together, our results suggest that the iDDR motif of TRF2 is the only region in the core telomeric complex capable of interacting with RAD50.

**Figure 2.**
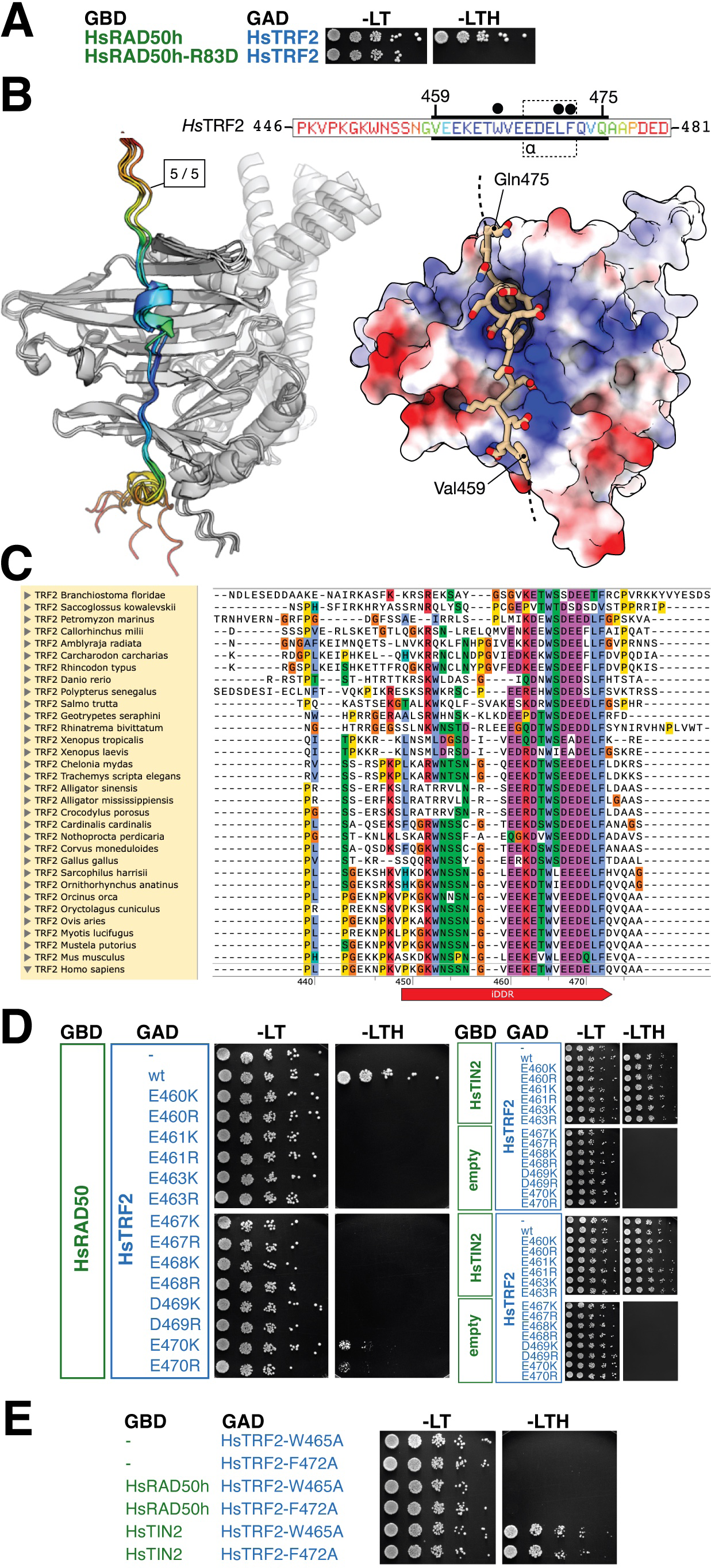
A model for binding of the iDDR motif to RAD50. (**A**) Yeast two-hybrid analysis of full-length HsTRF2 fused to the GAD domain of Gal4 for interaction against GBD-HsRAD50 encompassing HsRAD50 amino acids (aa) 1-221,1098-1312 either in wild-type form or carrying a R83D mutation. (**B**) Left: superposition of five independent AlphaFold2 models for the head domain of human RAD50 in complex with the iDDR of HsTRF2. Models are shown in cartoon representation, where HsRAD50 is coloured in grey and the identified interacting region of HsTRF2 coloured according to pLDDT score, using a continuous ‘rainbow’ spectrum from red (low confidence) to blue (high confidence). The amino sequence for the displayed section of HsTRF2 (aa 446-481) is provided as an inset. The thick black bars indicate the portion of sequence included in the model displayed on the right. An interacting region of HsTRF2 comprising amino acids Gly458-Gln475 was predicted with high confidence and consistency, containing a short helix segment spanning amino acids 468-472 (see inset: indicated with a dotted outline and labelled ‘ɑ’). Right: molecular surface of HsRAD50 coloured with respect to electrostatic potential. The top-ranked model for the interacting region of HsTRF2 is shown in stick-representation, with backbone carbon atoms coloured in ‘light tan’. This visualisation reveals a high degree of charge complementarity between the two interacting proteins. Amino acids residues Trp465, Leu471 and Phe472 of TRF2 coalesce to form a hydrophobic ‘plug’ that inserts itself into a small receiving pocket found on the surface of the HsRAD50 head domain (indicated by filled black circles above the primary sequence). (**C**) Multiple amino acid sequence alignment for putative iDDR regions from the indicated species, executed and displayed using the Clustal Omega functionality of SnapGene v.6. The annotation and numbering provided at the bottom of the alignment corresponds to the amino acid sequence of the human human protein. (**D**) and (**E**). Yeast two-hybrid analysis as in (A) of full-length HsTRF2 variants carrying the indicated mutations. All HsTRF2 variants were also tested against full-length HsTIN2 fusions to confirm protein functionality.

### The iDDR motif binds to the regulatory interface of RAD50 also recognised by the MIN motif

Previous analyses for binding of the yeast MIN motif to RAD50 pointed to an interaction with one surface of a β-sheet located in the N-terminal half of the globular head domain of RAD50 (Roisné-Hamelin et al., 2021; Marsella et al., 2021; Khayat et al., 2021). We refer to this as the ‘S’ region of RAD50, from the name previously given to the class of *S. cerevisiae* Rad50 alleles originally identified on the basis of their meiotic phenotype (Alani et al., 1990). The archetypal *rad50-S* allele is rad50-K81I, which results in loss of interaction with both Sae2 and the Rif2 MIN motif (Cannavo et al., 2018; Roisné-Hamelin et al., 2021; Marsella et al., 2021; Khayat et al., 2021). We therefore mutated the corresponding arginine residue in human RAD50 to aspartic acid and tested the ability of this mutant (R83D) to interact with TRF2. TRF2-R83D failed to bind RAD50 (Figure 2A) suggesting that the iDDR, like the MIN motif in yeast, binds to the S region of RAD50.

To test this idea further, we used the multimer mode of AlphaFold2 using ColabFold (Mirdita et al., 2022) to generate potential models for the iDDR motif in complex with RAD50. Each of the five resultant models had high confidence scores and a consistent mode of interaction (Figure 2B). The predicted RAD50-interacting region within the iDDR motif runs approximately from aa 459 to 475 of TRF2 and is arranged perpendicularly to the strands forming the β-sheet in RAD50. Reassuringly, Arg83 makes a specific contact with Asp469 of TRF2 in the model (see below, Figure 3B). Interestingly, aa 459-475 are also the most strongly conserved positions within the iDDR itself (Figure 2C). In the AlphaFold2-model, the conserved region also contains a run of charged amino acids positioned to interact with side-chains of RAD50 carrying an opposing charge (Figure 2B, right). We tested the relevance of these interactions in yeast two-hybrid assays, by use of several charge-swap mutations: each of the charged amino acid residues within TRF2 region 461-470 tested were found to be important for interaction, with only Glu470 retaining some residual activity in the binding assay (Figure 2D). We also confirmed the importance of the two single most conserved residues within the iDDR, Trp465 and Phe472: changes to alanine at either of these positions abrogated the ability of TRF2 to bind RAD50 (Figure 2E), in agreement with them coming together with Leu471 to form a hydrophobic bundle or ‘plug’ that interacts with a pocket on the surface of RAD50 (see below).

**Figure 3.**
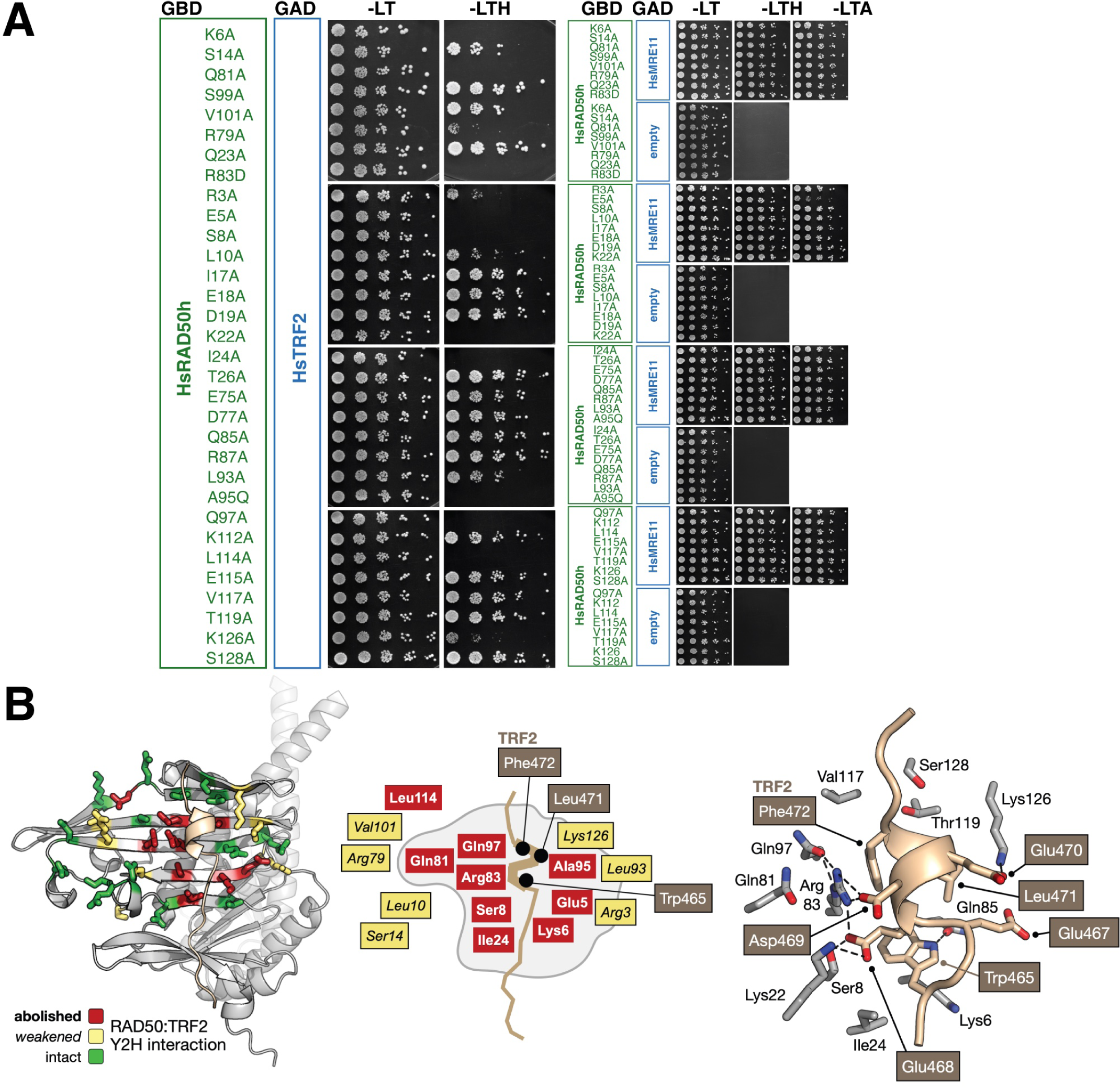
Map of RAD50 binding to iDDR. (**A**) Yeast two-hybrid analysis of full-length HsTRF2 fused to the GAD domain of Gal4 for interaction against GBD-HsRAD50 encompassing HsRAD50 amino acids 1-221,1098-1312 either in wild-type form or carrying the indicated mutations. In all cases the HsRAD50 variants were tested against full-length HsMRE11 to confirm protein functionality, and with empty vector to rule out self-activation. (**B**) Left: mapping of yeast two-hybrid data onto the surface of HsRAD50. The side chains for each mutation tested are shown in stick representation, coloured with respect to their effect on HsTRF2 interaction: red = interaction is abolished, yellow = interaction is weakened, green = interaction is intact / unaffected. Middle: schematic representation of the interactions mapped at the left, using the same colour scheme, with amino acids affecting the interaction with iDDR annotated. Right: schematic providing additional detail for the mapped yeast 2-hybrid data, showing amino acids in close proximity to the modelled TRF2-helix. Predicted hydrogen-bonds are represented by black dotted lines. Selected amino acids of HsTRF2 are annotated in white text on a grey/fawn background.

We further validated the binding model by mutating a large set of residues within the S region of RAD50, selecting those residues with side chains protruding towards the surface of the protein, and analysing their ability to interact with TRF2 by Y2H assay (Figure 3A). This analysis yielded data consistent with the AlphaFold2 model, with mutations reducing or ablating interaction mapping with close proximity to the position of iDDR in the model (Figure 3B, left and middle). The hydrophobic grouping of TRF2 residues Trp465, Leu471 and Phe472 is positioned directly above Ala95 of RAD50: a change of this residue to a bulkier glutamine side chain was found to disrupt binding. Additional mutations in the S region of RAD50, both to the left and to the right of the modelled C-terminal segment of iDDR, were also found to either ablate or partially reduce binding.

Taken together these results provide direct experimental support for a specific interaction of the iDDR motif with a regulatory region within RAD50, thus suggesting that TRF2 might employ a similar mechanism for MRN inactivation as that enforced by the MIN motif in yeast.

### The MIN motifs of Rif2 and Taz1 bind RAD50 at a position similar to that of the iDDR motif

Because the MIN and iDDR motifs lack any amino acid sequence similarity but share a common binding partner in RAD50 - which in eukaryotes is highly conserved in amino acid sequence particularly in the globular region - we also modelled binding of ScRif2 and SpTaz1 MIN motifs to their respective RAD50 partners, to assess any universality of this mechanism for MRN inactivation. AlphaFold2 predicted interactions in both cases; again, like for iDDR, the regions of high confidence in the models corresponded with the most conserved regions of both motifs (Figure 4A and B). In both the *S. cerevisiae* and *S. pombe* models, the N-terminal section of the MIN motif adopted an alpha helical structure, that bound in close proximity to the side chain of Rad50-Lys81. Pleasingly, this amino acid is required for interaction in both cases (Roisné-Hamelin et al., 2021; Marsella et al., 2021; Khayat et al., 2021; Cannavo et al., 2018). In addition, the ScRif2 model is remarkably consistent (Figure 4A) with a previous mutational analysis carried out for budding yeast RAD50 (Roisné-Hamelin et al., 2021). We note, however, that asparagine 18, located to the far left of the RAD50 β-sheet was also shown to affect interaction with Rif2 when mutated to serine (Marsella et al., 2021). We speculate that this mutation might affect interaction with the more C-terminal section of the Rif2 MIN motif, not accounted for in our model, but required for binding in yeast two-hybrid assays (Roisné-Hamelin et al., 2021). For the two MIN models, there is a remarkable degree of similarity in the interactions made by the central part of each motif, with Phe8/Phe511 of RIF2/Taz1 respectively, sitting in a small pocket on the surface of Rad50, along with identical positions of residues Pro10/Pro513, and Arg13/Arg515 as part of a larger phenylalanine-hydrophobic-proline-hydrophobic-arginine (FφPφR) motif (Figure 4A and B). In both cases, a short, predominantly negatively-charged helical element precedes the FφPφR motif, contributing to the binding interface through interaction with a reciprocal, positively charged Rad50 surface. The electrostatic charge distribution of the S regions in the two yeasts is, however, quite different across the lower part of each Rad50 molecule (as illustrated in Figure 4), with ScRad50 having a more pronounced negatively charged surface, which matches the propensity of the ScMIN motif to bear positively charged residues near the C-terminal end of the minimal sequence (Figure 4A) (Khayat et al., 2021), whereas the SpMIN at the equivalent positions contains negatively charged resides (Figure 4B).

**Figure 4.**
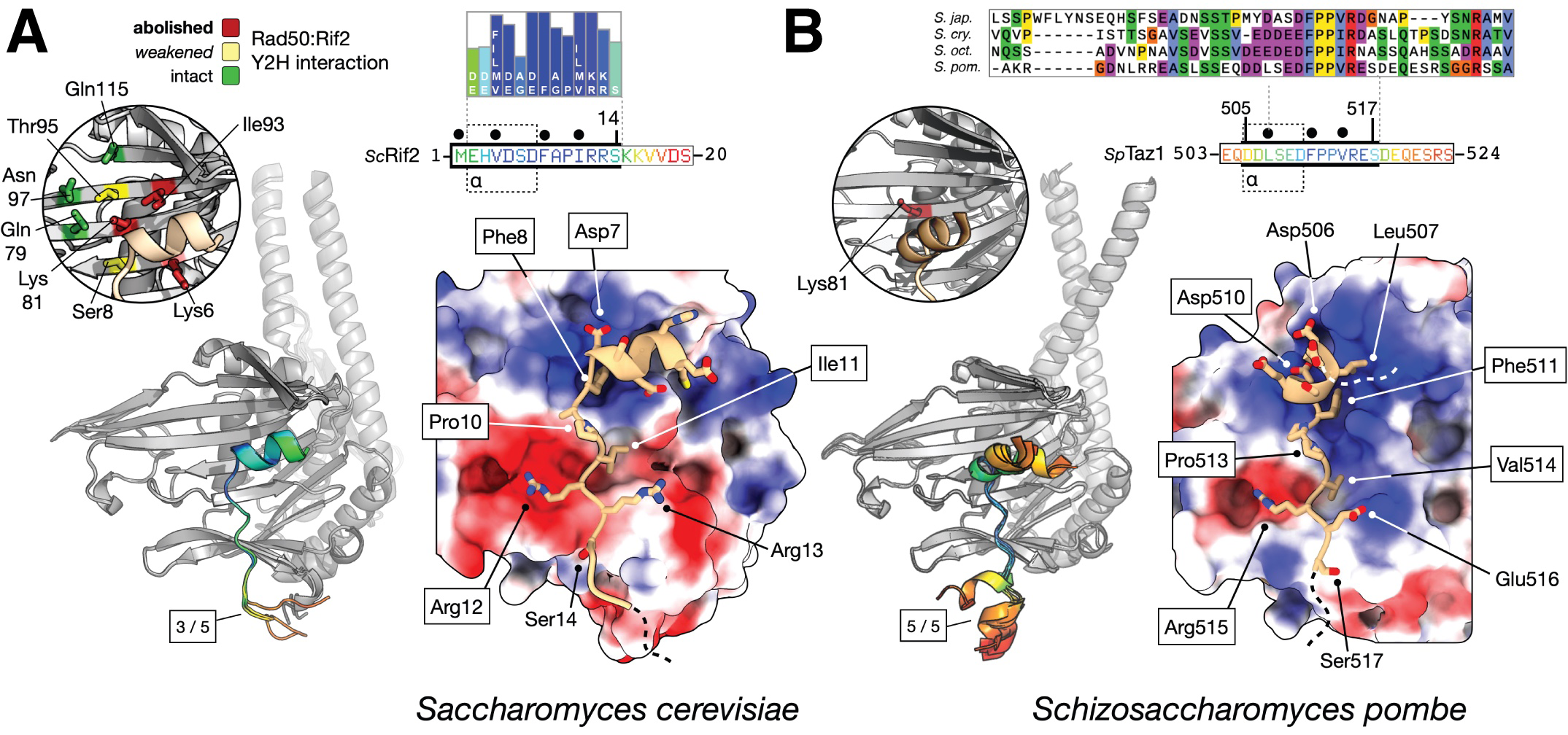
Models for ScRif2 and SpTaz1 MIN interactions with Rad50. (**A**) Left: superposition of three independent AlphaFold2 models for the head domain of ScRad50 in complex with the MIN motif of ScRif2. Models are shown in cartoon representation, where ScRad50 is coloured in grey and the identified interacting region of ScRif2 coloured according to pLDDT score using a continuous ‘rainbow’ spectrum from red (low confidence) to blue (high confidence. The primary sequence for the first 20 amino acids of ScRif2, including the MIN motif, is provided as an inset, using the same pLDDT colour scheme and aligned to the MIN consensus derived from Orc4 and Rif2 sequences (Khayat et al., 2021). Thick black bars indicate the portion of sequence included in the model displayed on the right. A region comprising amino acids 1-15 was predicted with high confidence and consistency, containing a short helix segment spanning amino acids 2-7 (see inset: indicated with a dotted outline and labelled ‘ɑ’). The circular inset, shown on the left maps the results of mutational analysis examining binding of ScRif2 to ScRad50 coloured as per Figure 3B (Roisné-Hamelin et al., 2021). Right: molecular surface of ScRad50 coloured with respect to electrostatic potential. The top-ranked model for the interacting region of ScRif2 is shown in stick-representation, with backbone carbon atoms coloured in ‘light tan’. This visualisation reveals a high degree of charge complementarity between the two interacting proteins. Amino acids of ScRif2, including Phe8, that make fully buried hydrophobic interactions with the surface of ScRad50 are annotated above the primary sequence inset, as shown by filled black circles. The five core positions in the MIN consensus from the Saccharomycotina [ScRif2 7-12 and SpTaz1 508-513] which allowed identification of MIN in SpTaz1 by homology search (Khayat et al., 2021) are bounded by a black rectangle in the annotation in both panels. (**B**) Predicted interaction between the SpTaz1 MIN motif and SpRad50, using the same representations as those shown in panel (A). A Clustal Omega alignment of the four Schizossacharomyces Taz1 amino acid sequences is provided as an inset. A region spanning amino acids 505-517 was predicted with high confidence and consistency, containing a short helix segment comprising amino acids 506-510 (see inset: indicated with a dotted outline and labelled ‘ɑ’). The secondary inset, highlights the location of Lys81, whose mutation prevents binding of SpTaz1 to SpRad50.

In conclusion, our structural predictions for each RAD50-MIN interaction are consistent with published mutational analyses that assess binding capability, and display an overall similarity to the predicted binding position of iDDR on RAD50, suggesting that these telomeric factors exploit a common mechanism to inactivate MRN and pointing to a shared mechanism of action.

### Binding of the iDDR/MIN motifs to RAD50 interferes with its ability to bind CtIP/Sae2/Ctp1

It was previously shown that the MIN motif of Rif2 impairs the endonucleolytic activity of MRN *in vitro,* as well as resection near DSBs when the MIN motif is tethered near a break site (Khayat et al., 2021; Roisné-Hamelin et al., 2021). In addition, Rif2 was shown to partly suppress the DNA repair defect of *sae2* mutants (Marsella et al., 2021). This has lead to the idea that the MIN motif might act to inhibit MRN nuclease action by preventing the association of the CtIP/ Sae2/Ctp1 cofactor (Marsella et al., 2021), which is required for DNA cleavage (Cannavo and Cejka, 2014; Anand et al., 2016; Deshpande et al., 2016). We therefore decided to investigate the binding mode of the three orthologues CtIP, Sae2 and Ctp1 to their respective RAD50 counterparts using AlphaFold2 as before. This analysis predicted interaction with the S region of RAD50 for all three proteins utilising the conserved N-terminal ‘Sae2-like’ region (Andres and Williams, 2017)(Figure 4A-C). Pleasingly, the model for ScSae2 agrees with the available genetic data and the distribution of the original *rad50-S* alleles (Figure 5B) (Alani et al., 1990; Roisné-Hamelin et al., 2021).

**Figure 5.**
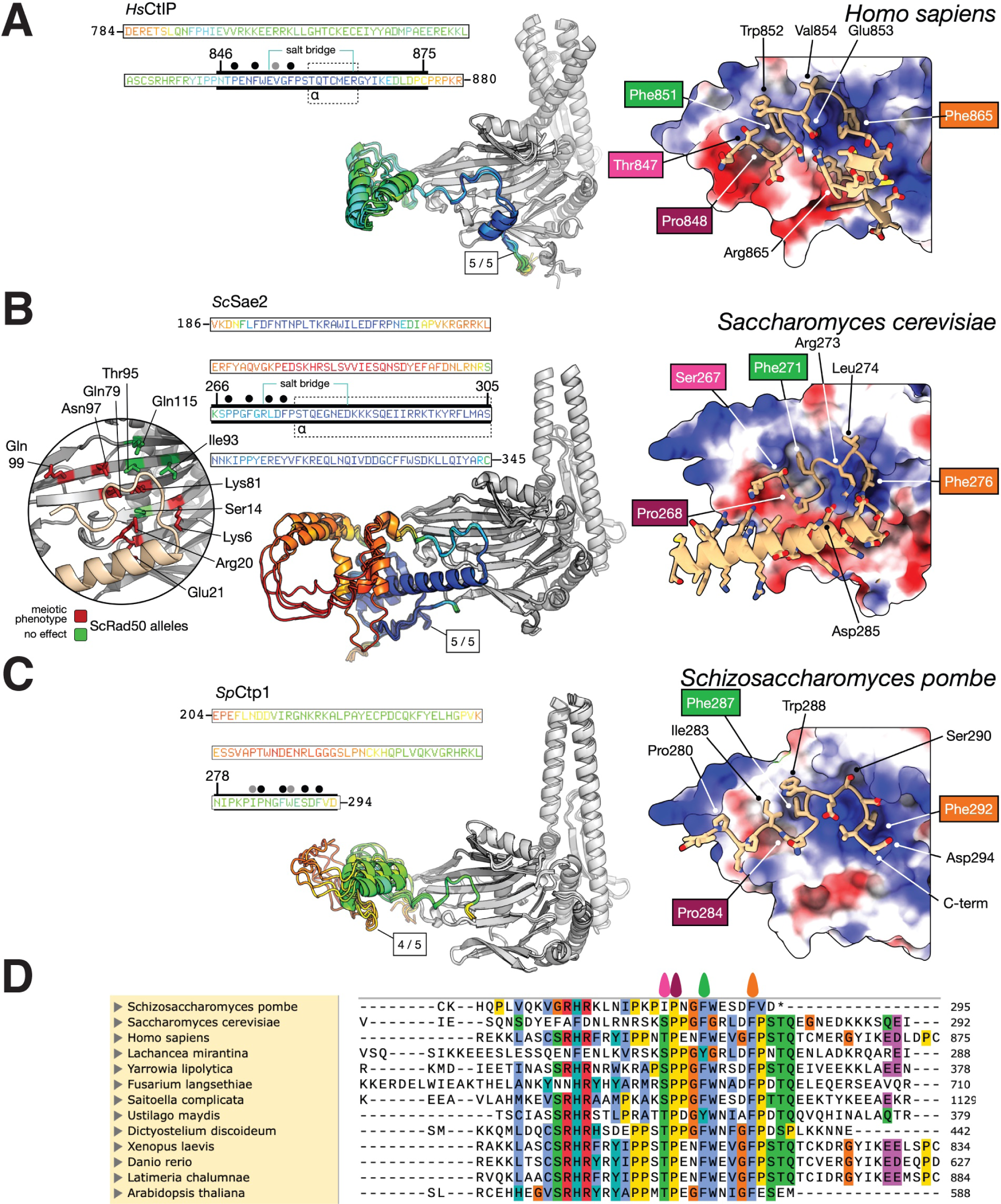
Models for HsCtIP, ScSae2 and SpCtp1 interactions with Rad50. Panels (**A**, **B** and **C**). Left: the number of AlphaFold2 models (out of the 5 generated) with consistent binding pose are indicated, shown in cartoon representation, and coloured according to pLDDT score using a continuous ‘rainbow’ spectrum from red (low confidence) to blue (high confidence). The head domain of RAD50 is also shown in cartoon representation and coloured grey. In each case, the amino acid sequence for the region predicted with high-to-medium confidence is provided as an inset. Thick black bars indicate the portion of sequence included in the figure displayed to the right. Right: Molecular surface of RAD50 coloured according to electrostatic potential, with the top-ranked model shown in combined cartoon / stick representation, and backbone carbon atoms coloured in ‘light tan’. (**A**) HsRAD50 in complex with CtIP. A region comprising amino acids Gln791-Asp873 was predicted with high confidence and consistency, containing a short helix segment spanning amino acids Thr859-Arg865 (see inset: indicated with a dotted outline and labelled ‘ɑ’). The model also suggests formation of an internal HsCtIP salt-bridge between the side chains of Glu853 and Arg865. Right: This visualisation reveals a high degree of charge complementarity between the two interacting proteins, with HsCTIP residues Pro848, Phe851, Val854 and Phe856 making additional hydrophobic interactions with the surface of the HsRAD50 head domain (highlighted by black filled circles placed above the inset primary sequence if fully buried, or dark grey if packed against the surface). Selected amino acid side chains are shown in stick representation, with labels coloured according to the ‘teardrops’ placed above the multiple amino acid sequence alignment shown in panel (D) for a few conserved residues, including HsCTIP-Thr847 as a known site of phosphorylation. (**B**) ScRad50 in complex with ScSae2. A region spanning amino acids Ser265-Cys345 was predicted with high confidence and consistency, containing a helix segment spanning amino acids Lys266-Ser305 (see inset: indicated with a dotted outline and labelled ‘ɑ’). The model also suggests formation of an internal ScSae2 salt-bridge between the side chains of Arg273 and Asp285. Residues Pro268, Phe271, Leu274 and Phe276 of ScSae2 make buried hydrophobic interactions with the surface of the ScRad50 head domain (highlighted by black filled circles placed above the inset primary sequence). The circular inset summarises genetic data for the possible interaction of ScSae2 with ScRad50 (Roisné-Hamelin et al., 2021; Alani et al., 1990). Right: selected amino acid side chains are shown in stick representation, with labels coloured according to the ‘teardrops’ placed above the multiple amino acid sequence alignment shown in panel (D) for a few conserved residues, including ScSae2-Ser267 as a known site of phosphorylation. (**C**) SpRad50 in complex with SpCtp1. A region spanning amino acids Gln265-Val293 was predicted with medium confidence and consistency. Residues Ile283, Pro284, Phe287, Trp288, Ser290 and Phe292 of SpCtp1 make hydrophobic interactions with the surface of the SpRad50 head domain (highlighted by black filled circles placed above the inset primary sequence if fully buried, or dark grey if packed against the surface). Right: selected amino acid side chains are shown in stick representation, with labels coloured according to the ‘teardrops’ placed above the multiple amino acid sequence alignment shown in panel (D) for a few conserved residues. (**D**) Multiple amino acid sequence alignment for the C-terminal region of HsCtIP/ScSae2/SpCtp1, plus homologues from the indicated species, executed and displayed using the Clustal Omega functionality of SnapGene v.6.

Here, the models identify two main areas, or ‘pockets’, for CtIP/Sae2/Ctp1 interaction with RAD50 (Figure 6A): Pocket A largely overlaps with the binding site for iDDR/MIN describe above, whereas Pocket B is located to its left (as represented in Figure 6A) in a part of the β-sheet region where our modelling did not identify strong iDDR/MIN binding. The predictions uncover similarities in the binding mode of CtIP/Sae2/Ctp1 to RAD50, placing recognised critical phosphorylation sites in Sae2 and CtIP (Ser267 and Thr847, respectively) for interaction in Pocket B (Huertas et al., 2008; Huertas and Jackson, 2009; Anand et al., 2016; Cannavo et al., 2018). Furthermore, the RAD50-interacting portion of SpCtp1 is restricted to the last 17 aa in the protein and almost exactly coincides with a C-terminal 15-aa peptide that is capable of stimulating MRN nucleolytic activity *in vitro* (Zdravković et al., 2021). Notably, the MIN and iDDR motifs and the Sae2-like domains all bear hydrophobic amino acids which are among the most highly invariant in each motif/domain and which make deep contacts into a conserved small cavity in RAD50: these are Phe8 in the ScMIN; Phe511 in the SpMIN; Trp465, Leu471 and Phe472 in the HsiDDR; Phe865, 276 and 292 in the HsCtIP, ScSae2 and SpCtp1 Sae2-like domains, respectively (Figures 2B, 3B, 4 and 5).

**Figure 6.**
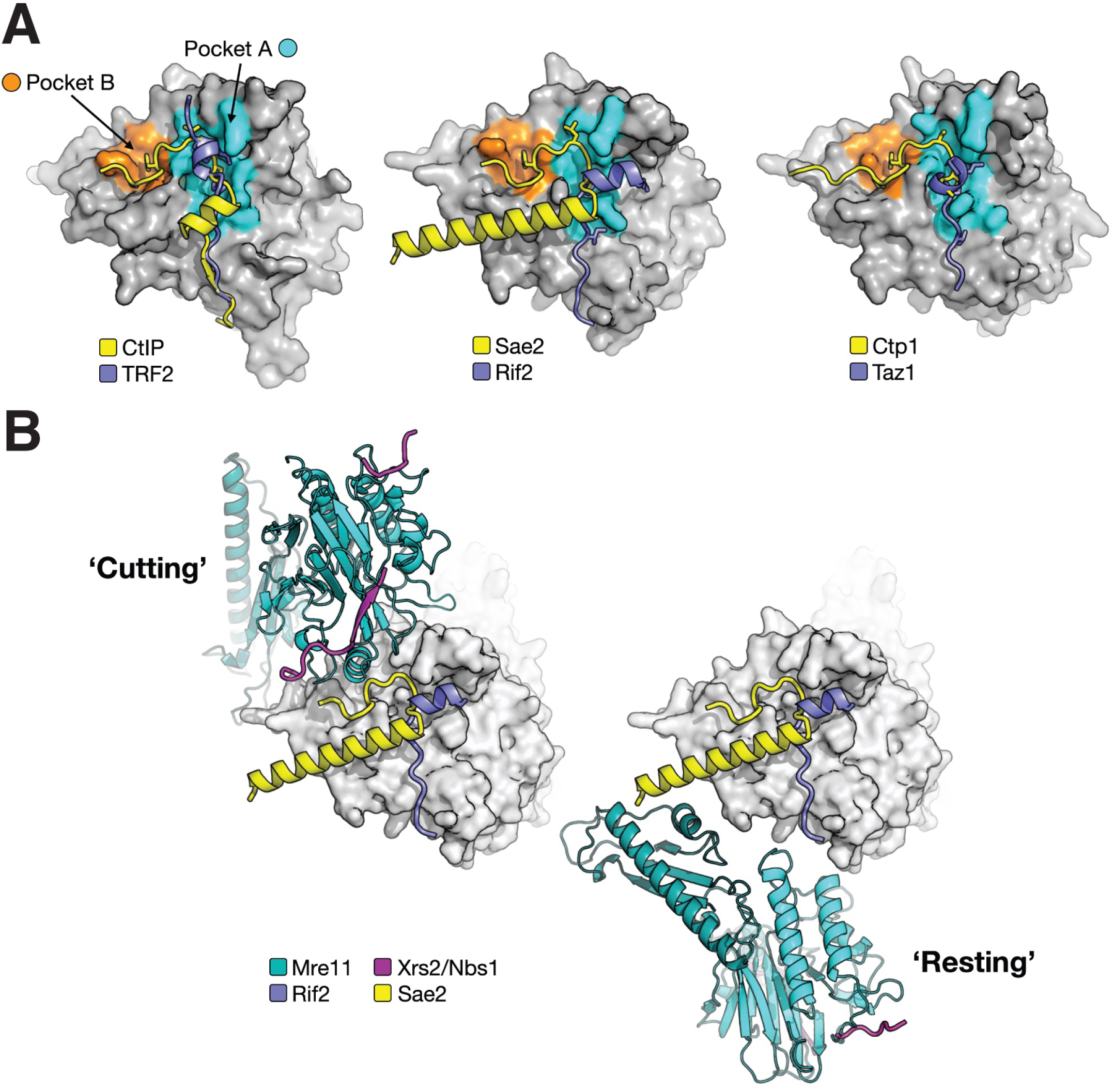
The iDDR/MIN and HsCtIP/ScSae2/SpCtp1 binding sites partially overlap. (**A**) Schematic summary of AlphaFold2-modelling for the interaction of each indicated motif, with the surface of RAD50 (coloured grey). iDDR/MIN motif predominantly interact with a singular pocket found on the surface of RAD50 (coloured cyan, and labelled ‘A’). The CtIP orthologues, however interact with both Pocket ‘A’ and a second pocket (coloured orange, and labelled ‘B’). (**B**) Presumed compatibility of modelled/mapped interactions with the ‘Cutting’ or ‘Resting’ states of the MRN/MRX complex. Models were generated by superposition of ScRad50-Sae2 and ScRad50-Sae2 models on the deposited cryoEM structures of the *E. Coli* Mre11-Rad50 (SbcCD) head complex (PDB IDs: 6S85 and 6S6V) (Käshammer et al., 2019). The expected location for Xrs2/Nbs1 binding to Mre11 was mapped through superposition of the X-ray crystal structure of fission yeast Mre11 in complex with a fragment of Nbs1 (PDB: 4FBW) (Schiller et al., 2012). Please see associated key for additional detail.

Our Y2H-data, plus that of others, combined with interaction modelling by AlphaFold2, reveal a pattern of interactions that identify a conserved mode of interaction for MRN co-factors in binding to RAD50, which is consistent with the notion that iDDR or MIN binding is mutually exclusive to that of CtIP/Sae2/Ctp1.

## DISCUSSION

Our previous work in budding yeast uncovered a conserved amino acid motif dedicated to inhibition of MRN activity, which first arose in DNA replication origin factor Orc4 and then, through sub-functionalisation, was retained by telomeric protein Rif2 (Khayat et al., 2021). We called this motif ‘MIN’, for MRN inhibitor, to highlight its potency in disabling the multiple facets of MRN complex activity. Here we provide structural modelling, obtained using AlphaFold2, that provides details for the most conserved section of the Rif2-RAD50 interaction, and which is consistent with previous mutational data examining their interaction (Roisné-Hamelin et al., 2021). Strikigly, a model for interaction of the MIN motif from SpTaz1 with SpRad50 predicts a similar set of interactions. In particular, several of the contacts made are highly similar between the two organisms, and closely correspond to the most conserved amino acids residues in a ‘pile-up’ compiled from Orc4 and Rif2 MIN motifs in the Saccharomycotina (Khayat et al., 2021), explaining why we were previously able to identify the MIN motif in the *Schizosaccharomyces* species based on amino acid sequence identity alone. However, changes in the electrostatic surface potential of the RAD50 S region, which are apparent in the three structures presented here, would by necessity lead to changes in the amino acid sequence composition of a MIN-like peptide, and therefore similar short RAD50-binding motifs/activities might be easily missed in analyses by protein sequence alone. For example, the MIN-motifs of Rif2 and Taz1 share a general preponderance of negatively charged amino acids at their respective N-terminal ends, whereas the region more C-terminal to the core of the motif contains a swap in the charge of a single amino acid, i.e. the equivalent of Rif2-Arg13 is Taz1-Glu516, thus matching a concomitant change in electrostatic potential on the surface of Rad50.

The combined genetic and biochemical analysis of Sae2 binding to Rad50 (Alani et al., 1990; Roisné-Hamelin et al., 2021; Cannavo et al., 2018) pointed at the potential of this RAD50 β-sheet region to bind MRN regulators. The data we present here demonstrate that the iDDR motif of human TRF2 makes use of the same region when binding to RAD50. The evolution of telomere DNA binding proteins appears complex and is not well understood, but TRF2 is possibly not a direct orthologue of Taz1, given that the two proteins are structurally different (Červenák et al., 2017; Myler et al., 2021; Deng et al., 2015; Lue, 2021). Regardless of any common ancestry, our analyses strongly indicates that the binding of the TRF2-iDDR motif to RAD50 represents a third example of evolution producing a RAD50 binding motif within a telomeric protein. The binding-position of both iDDR and MIN motifs on the surface of RAD50 is highly similar, in both cases containing a short alpha-helix that binds at the same location on Rad50, flanked by a patch of amino acids complementary to the charge present on the corresponding interaction surface of RAD50 (compare Figures 2B and 4A,B). However, the position of the charged stretch of amino acids is reversed in the two systems, with the alpha helix sitting at its C-terminal side in iDDR, but at the N-terminal side in the two MIN motifs. While in principle the different distribution of the charged amino acids in the iDDR and MIN could have diverged to match the electrostatic potential of RAD50, the observed inversion in polypeptide strand polarity, together with the difference in hydrophobic interactions - iDDR forming a distinctive hydrophobic ‘plug’ comprising amino acids Trp465, Leu471 and Phe472 - demonstrate the that the iDDR/RAD50 interaction constitutes a third example of convergent evolution for this particular mechanism of RAD50 binding.

The models presented here for CtIP/Sae2/Ctp1 binding to RAD50 serve to validate earlier suggestions that the MIN motif might prevent endonucleolytic action by MRN by directly interfering with ability of the required CtIP/Sae2/Ctp1 co-factor to bind RAD50 (Roisné-Hamelin et al., 2021; Marsella et al., 2021; Khayat et al., 2021; Cannavo et al., 2018). Our analysis has identified two main binding ‘pockets’ for CtIP/Sae2/Ctp1 binding to RAD50, one of which fully overlaps the iDDR/MIN binding site, thus confirming that CtIP/Sae2/Ctp1 and iDDR/MIN binding are mutually exclusive (Bonetti et al., 2021). However, as parts of the extended MIN and iDDR motifs are only poorly modelled by AlphaFold2, it remains possible that the overlap in binding is even more extensive. These findings present a scenario of opposing regulatory activities exemplified by a positive regulator of MRN in the CtIP/Sae2/Ctp1 orthologues and a set of negative regulators in telomeric factors Rif2, Taz1 and TRF2. The interplay between these regulatory factors is, however, not clear and the relative affinities of each protein or interaction motif for RAD50 remain to be determined. However, due to its hydrophobic plug and thus larger buried surface area, the iDDR motif in particular might be able to bind RAD50 with high affinity. In addition, the local high concentration of telomeric factors are likely to shift the balance in favour of MRN inhibition specifically at chromosome ends.

ATP binding by RAD50 produces an allosteric switch in the MRN complex that suppresses endonuclease action, but is proficient for both DNA binding and kinase activation (Hailemariam et al., 2019b; Deshpande et al., 2014; Lafrance-Vanasse et al., 2015). The hydrolysis of ATP results in a repositioning of the MRE11 dimer from a position siting ‘below’ the RAD50 head-domain in the ‘resting’ state, to a ‘up’ or ‘sideways’ position compared to RAD50 in the ‘cutting’ conformation (Figure 6B) (Gut et al., 2022; Käshammer et al., 2019). Binding of CtIP/Sae2/Ctp1 has been suggested to promote the ‘cutting’ conformation by stabilising an MRE11-RAD50 interaction (Figure 6B, left) (Käshammer et al., 2019). The proximity and complementarity of the modelled Sae2 interaction with Mre11, suggests that it helps to form a composite interaction site, promoting the cutting state, in the absence of the ‘fastener’ loop present in the bacterial orthologues of Mre11 (Marsella et al., 2021; Käshammer et al., 2019).

As discussed above, the iDDR/MIN motifs are expected to prevent CtIP/Sae2/Ctp1 binding, and therefore stabilisation of the ‘cutting’ state of MRN. In budding yeast, Rif2 has been shown to increase the ATPase activity of RAD50 through the action of the MIN motif and this particular property has been invoked as being instrumental in mediating the effect of Rif2 in suppressing NHEJ and ATM/Tel1 activation at telomeres (Cassani et al., 2016; Hailemariam et al., 2019a; Khayat et al., 2021; Roisné-Hamelin et al., 2021). While this scenario is attractive, it is presently not clear whether the iDDR motif might have a similar effect on human MRN, nor do the models presented here offer a clear insight into how the MIN/iDDR motif might affect ATP hydrolysis by RAD50. Further work will be required to fully elucidate the precise role of the iDDR and MIN motifs in the modulation of MRN activity at telomeres, but the analyses that we present here underscore how this is a ‘problem’ that has been addressed multiple times at telomeres during evolution, with a similar solution.

## ACKNOWLEDGMENTS

AB wishes to express gratitude to a group of enthusiastic undergraduate students who helped make and test a number of RAD50 and TRF2 mutations: Tom Blount, Amy Cheung, Phoebe Frewer, Sabiha Kahn, Joe Kleinschmidt, Thomas Panzar, Rosie Russell, Kataharine Stutz, Hannah Taylor.

## FUNDING

This work was supported by Cancer Research UK [C302/A24386 to AWO and Prof. Laurence H. Pearl, University of Sussex]. MA acknowledges support from the Saudi Arabian Cultural Bureau.

## Conflict of interest statement

none declared.

## Notes

### Competing Interest Statement

The authors have declared no competing interest.

### Summary of Updates

The new version uses a different negative control in Figure 1C (a point mutation in iDDR that abolishes the interaction with RAD50), and makes minor corrections in the text and in the figures (for example it adjusts the positioning of the teardrops on top of the Clustal alignment in Figure 5D).

## REFERENCES

Alani, E., Padmore, R., and Kleckner, N. (1990). Analysis of wild-type and rad50 mutants of yeast suggests an intimate relationship between meiotic chromosome synapsis and recombination. Cell 61, 419–436.

Anand, R., Ranjha, L., Cannavo, E., and Cejka, P. (2016). Phosphorylated CtIP Functions as a Co-factor of the MRE11-RAD50-NBS1 Endonuclease in DNA End Resection. Mol Cell

Andres, S. N., and Williams, R. S. (2017). CtIP/Ctp1/Sae2, molecular form fit for function. DNA Repair (Amst) 56, 109–117.

Attwooll, C. L., Akpinar, M., and Petrini, J. H. (2009). The mre11 complex and the response to dysfunctional telomeres. Mol Cell Biol 29, 5540–5551.

Bianchi, A., and Shore, D. (2007). Increased association of telomerase with short telomeres in yeast. Genes Dev 21, 1726–1730.

Bonetti, D., Clerici, M., and Longhese, M. P. (2021). Interplay between Sae2 and Rif2 in the regulation of Mre11-Rad50 activities at DNA ends. Curr Opin Genet Dev 71, 72–77.

Boulton, S. J., and Jackson, S. P. (1998). Components of the Ku-dependent non-homologous end-joining pathway are involved in telomeric length maintenance and telomeric silencing. Embo J 17, 1819–1828.

Cannavo, E., Johnson, D., Andres, S. N., Kissling, V. M., Reinert, J. K., Garcia, V., Erie, D. A., Hess, D., Thomä, N. H., Enchev, R. I., Peter, M., Williams, R. S., Neale, M. J., and Cejka, P. (2018). Regulatory control of DNA end resection by Sae2 phosphorylation. Nat Commun 9, 4016.

Cannavo, E., and Cejka, P. (2014). Sae2 promotes dsDNA endonuclease activity within Mre11-Rad50-Xrs2 to resect DNA breaks. Nature 514, 122–125.

Cassani, C., Gobbini, E., Wang, W., Niu, H., Clerici, M., Sung, P., and Longhese, M. P. (2016). Tel1 and Rif2 Regulate MRX Functions in End-Tethering and Repair of DNA Double-Strand Breaks. PLoS Biol 14, e1002387.

Červenák, F., Juríková, K., Sepšiová, R., Neboháčová, M., Nosek, J., and Tomáška, L. (2017). Double-stranded telomeric DNA binding proteins: Diversity matters. Cell Cycle 16, 1568–1577.

Červenák, F., Sepšiová, R., Nosek, J., and Tomáška, Ľ. (2021). Step-by-Step Evolution of Telomeres: Lessons from Yeasts. Genome Biol Evol 13, evaa268.

Craven, R. J., and Petes, T. D. (1999). Dependence of the regulation of telomere length on the type of subtelomeric repeat in the yeast Saccharomyces cerevisiae. Genetics 152, 1531–1541.

de Lange, T. (2018). Shelterin-Mediated Telomere Protection. Annu Rev Genet 52, 223–247.

Deng, W., Wu, J., Wang, F., Kanoh, J., Dehe, P.-M., Inoue, H., Chen, J., and Lei, M. (2015). Fission yeast telomere-binding protein Taz1 is a functional but not a structural counterpart of human TRF1 and TRF2. Cell Res 25, 881–884.

Deng, Y., Guo, X., Ferguson, D. O., and Chang, S. (2009). Multiple roles for MRE11 at uncapped telomeres. Nature 460, 914–918.

Deshpande, R. A., Lee, J. H., Arora, S., and Paull, T. T. (2016). Nbs1 Converts the Human Mre11/ Rad50 Nuclease Complex into an Endo/Exonuclease Machine Specific for Protein-DNA Adducts. Mol Cell 64, 593–606.

Deshpande, R. A., Williams, G. J., Limbo, O., Williams, R. S., Kuhnlein, J., Lee, J.-H., Classen, S., Guenther, G., Russell, P., Tainer, J. A., and Paull, T. T. (2014). ATP-driven Rad50 conformations regulate DNA tethering, end resection, and ATM checkpoint signaling. EMBO J 33, 482–500.

Dimitrova, N., and de Lange, T. (2009). Cell cycle-dependent role of MRN at dysfunctional telomeres: ATM signaling-dependent induction of nonhomologous end joining (NHEJ) in G1 and resection-mediated inhibition of NHEJ in G2. Mol Cell Biol 29, 5552–5563.

Faure, V., Coulon, S., Hardy, J., and Geli, V. (2010). Cdc13 and telomerase bind through different mechanisms at the lagging- and leading-strand telomeres. Mol Cell 38, 842–852.

Gibson, D. G. (2011). Enzymatic assembly of overlapping DNA fragments. Methods Enzymol 498, 349–361.

Gut, F., Käshammer, L., Lammens, K., Bartho, J. D., Boggusch, A. M., van de Logt, E., Kessler, B., and Hopfner, K. P. (2022). Structural mechanism of endonucleolytic processing of blocked DNA ends and hairpins by Mre11-Rad50. Mol Cell 82, 3513–3522.e6.

Hailemariam, S., De Bona, P., Galletto, R., Hohl, M., Petrini, J. H., and Burgers, P. M. (2019a). The telomere-binding protein Rif2 and ATP-bound Rad50 have opposing roles in the activation of yeast Tel1^ATM^ kinase. J Biol Chem 294, 18846–18852.

Hailemariam, S., Kumar, S., and Burgers, P. M. (2019b). Activation of Tel1^ATM^ kinase requires Rad50 ATPase and long nucleosome-free DNA, but no DNA ends. J Biol Chem

Hector, R. E., Shtofman, R. L., Ray, A., Chen, B. R., Nyun, T., Berkner, K. L., and Runge, K. W. (2007). Tel1p preferentially associates with short telomeres to stimulate their elongation. Mol Cell 27, 851–858.

Hopfner, K. P. (2023). Mre11-Rad50: the DNA end game. Biochem Soc Trans BST20220754.

Huertas, P., and Jackson, S. P. (2009). Human CtIP Mediates Cell Cycle Control of DNA End Resection and Double Strand Break Repair. J Biol Chem 284, 9558–9565.

Huertas, P., Cortés-Ledesma, F., Sartori, A. A., Aguilera, A., and Jackson, S. P. (2008). CDK targets Sae2 to control DNA-end resection and homologous recombination. Nature 455, 689–692.

Kaizer, H., Connelly, C. J., Bettridge, K., Viggiani, C., and Greider, C. W. (2015). Regulation of Telomere Length Requires a Conserved N-Terminal Domain of Rif2 in Saccharomyces cerevisiae. Genetics

Karlseder, J., Hoke, K., Mirzoeva, O. K., Bakkenist, C., Kastan, M. B., Petrini, J. H., and Lange de, T. (2004). The Telomeric Protein TRF2 Binds the ATM Kinase and Can Inhibit the ATM-Dependent DNA Damage Response. PLoS Biol 2, E240.

Käshammer, L., Saathoff, J. H., Lammens, K., Gut, F., Bartho, J., Alt, A., Kessler, B., and Hopfner, K. P. (2019). Mechanism of DNA End Sensing and Processing by the Mre11-Rad50 Complex. Mol Cell

Khayat, F., Cannavo, E., Alshmery, M., Foster, W. R., Chahwan, C., Maddalena, M., Smith, C., Oliver, A. W., Watson, A. T., Carr, A. M., Cejka, P., and Bianchi, A. (2021). Inhibition of MRN activity by a telomere protein motif. Nat Commun 12, 3856.

Lafrance-Vanasse, J., Williams, G. J., and Tainer, J. A. (2015). Envisioning the dynamics and flexibility of Mre11-Rad50-Nbs1 complex to decipher its roles in DNA replication and repair. Prog Biophys Mol Biol 117, 182–193.

Lazzerini-Denchi, E., and Sfeir, A. (2016). Stop pulling my strings - what telomeres taught us about the DNA damage response. Nat Rev Mol Cell Biol 17, 364–378.

Lee, J.-H., Mand, M. R., Deshpande, R. A., Kinoshita, E., Yang, S.-H., Wyman, C., and Paull, T. T. (2013). Ataxia telangiectasia-mutated (ATM) kinase activity is regulated by ATP-driven conformational changes in the Mre11/Rad50/Nbs1 (MRN) complex. J Biol Chem 288, 12840–12851.

Lee, S. S., Bohrson, C., Pike, A. M., Wheelan, S. J., and Greider, C. W. (2015). ATM Kinase Is Required for Telomere Elongation in Mouse and Human Cells. Cell Rep 13, 1623–1632.

Lim, C. J., and Cech, T. R. (2021). Shaping human telomeres: from shelterin and CST complexes to telomeric chromatin organization. Nat Rev Mol Cell Biol

Lue, N. F. (2021). Duplex Telomere-Binding Proteins in Fungi With Canonical Telomere Repeats: New Lessons in the Rapid Evolution of Telomere Proteins. Front Genet 12, 638790.

Lustig, A. J. (2019). Towards the Mechanism of Yeast Telomere Dynamics. Trends Cell Biol 29, 361–370.

Marcand, S., Pardo, B., Gratias, A., Cahun, S., and Callebaut, I. (2008). Multiple pathways inhibit NHEJ at telomeres. Genes Dev 22, 1153–1158.

Marsella, A., Gobbini, E., Cassani, C., Tisi, R., Cannavo, E., Reginato, G., Cejka, P., and Longhese, M. P. (2021). Sae2 and Rif2 regulate MRX endonuclease activity at DNA double-strand breaks in opposite manners. Cell Rep 34, 108906.

McGee, J. S., Phillips, J. A., Chan, A., Sabourin, M., Paeschke, K., and Zakian, V. A. (2010). Reduced Rif2 and lack of Mec1 target short telomeres for elongation rather than double-strand break repair. Nature structural & molecular biology 17, 1438–1445.

Mirdita, M., Schütze, K., Moriwaki, Y., Heo, L., Ovchinnikov, S., and Steinegger, M. (2022). ColabFold: making protein folding accessible to all. Nat Methods 19, 679–682.

Myler, L. R., Kinzig, C. G., Sasi, N. K., Zakusilo, G., Cai, S. W., and de Lange, T. (2021). The evolution of metazoan shelterin. Genes Dev 35, 1625–1641.

Nugent, C. I., Bosco, G., Ross, L. O., Evans, S. K., Salinger, A. P., Moore, J. K., Haber, J. E., and Lundblad, V. (1998). Telomere maintenance is dependent on activities required for end repair of double-strand breaks. Curr Biol 8, 657–660.

Okamoto, K., Bartocci, C., Ouzounov, I., Diedrich, J. K., Yates, J. R., and Denchi, E. L. (2013). A two-step mechanism for TRF2-mediated chromosome-end protection. Nature 494, 502–505.

Park, Y. B., Hohl, M., Padjasek, M., Jeong, E., Jin, K. S., Krężel, A., Petrini, J. H., and Cho, Y. (2017). Eukaryotic Rad50 functions as a rod-shaped dimer. Nat Struct Mol Biol 24, 248–257.

Pettersen, E. F., Goddard, T. D., Huang, C. C., Meng, E. C., Couch, G. S., Croll, T. I., Morris, J. H., and Ferrin, T. E. (2021). UCSF ChimeraX: Structure visualization for researchers, educators, and developers. Protein Sci 30, 70–82.

Rai, R., Hu, C., Broton, C., Chen, Y., Lei, M., and Chang, S. (2017). NBS1 Phosphorylation Status Dictates Repair Choice of Dysfunctional Telomeres. Mol Cell 65, 801–817.e4.

Reis, C. C., Batista, S., and Ferreira, M. G. (2012). The fission yeast MRN complex tethers dysfunctional telomeres for NHEJ repair. EMBO J 31, 4576–4586.

Roisné-Hamelin, F., Pobiega, S., Jézéquel, K., Miron, S., Dépagne, J., Veaute, X., Busso, D., Du, M. L., Callebaut, I., Charbonnier, J. B., Cuniasse, P., Zinn-Justin, S., and Marcand, S. (2021). Mechanism of MRX inhibition by Rif2 at telomeres. Nat Commun 12, 2763.

Roset, R., Inagaki, A., Hohl, M., Brenet, F., Lafrance-Vanasse, J., Lange, J., Scandura, J. M., Tainer, J. A., Keeney, S., and Petrini, J. H. J. (2014). The Rad50 hook domain regulates DNA damage signaling and tumorigenesis. Genes Dev 28, 451–462.

Rotheneder, M., Stakyte, K., van de Logt, E., Bartho, J. D., Lammens, K., Fan, Y., Alt, A., Kessler, B., Jung, C., Roos, W. P., Steigenberger, B., and Hopfner, K. P. (2023). Cryo-EM structure of the Mre11-Rad50-Nbs1 complex reveals the molecular mechanism of scaffolding functions. Mol Cell 83, 167–185.e9.

Sabourin, M., Tuzon, C. T., and Zakian, V. A. (2007). Telomerase and Tel1p preferentially associate with short telomeres in S. cerevisiae. Mol Cell 27, 550–561.

Schiller, C. B., Lammens, K., Guerini, I., Coordes, B., Feldmann, H., Schlauderer, F., Möckel, C., Schele, A., Strässer, K., Jackson, S. P., and Hopfner, K.-P. (2012). Structure of Mre11-Nbs1 complex yields insights into ataxia-telangiectasia-like disease mutations and DNA damage signaling. Nat Struct Mol Biol 19, 693–700.

Schrödinger, L. L. C. The PyMOL Molecular Graphics System, Version 2.0. https://pymol.org/2

Soudet, J., Jolivet, P., and Teixeira, M. T. (2014). Elucidation of the DNA End-Replication Problem in Saccharomyces cerevisiae. Mol Cell 53, 954–964.

Syed, A., and Tainer, J. A. (2018). The MRE11-RAD50-NBS1 Complex Conducts the Orchestration of Damage Signaling and Outcomes to Stress in DNA Replication and Repair. Annu Rev Biochem 87, 263–294.

Tong, A. S., Stern, J. L., Sfeir, A., Kartawinata, M., de Lange, T., Zhu, X. D., and Bryan, T. M. (2015). ATM and ATR Signaling Regulate the Recruitment of Human Telomerase to Telomeres. Cell Rep 13, 1633–1646.

Viscardi, V., Baroni, E., Romano, M., Lucchini, G., and Longhese, M. P. (2003). Sudden Telomere Lengthening Triggers a Rad53-dependent Checkpoint in Saccharomyces cerevisiae. Mol Biol Cell 14, 3126–3143.

Wellinger, R. J., and Zakian, V. A. (2012). Everything you ever wanted to know about Saccharomyces cerevisiae telomeres: beginning to end. Genetics 191, 1073–1105.

Wu, P., van Overbeek, M., Rooney, S., and de Lange, T. (2010). Apollo contributes to G overhang maintenance and protects leading-end telomeres. Mol Cell 39, 606–617.

Xue, J., Chen, H., Wu, J., Takeuchi, M., Inoue, H., Liu, Y., Sun, H., Chen, Y., Kanoh, J., and Lei, M. (2017). Structure of the fission yeast S. pombe telomeric Tpz1-Poz1-Rap1 complex. Cell Res 27, 1503–1520.

Zdravković, A., Daley, J. M., Dutta, A., Niwa, T., Murayama, Y., Kanamaru, S., Ito, K., Maki, T., Argunhan, B., Takahashi, M., Tsubouchi, H., Sung, P., and Iwasaki, H. (2021). A conserved Ctp1/ CtIP C-terminal peptide stimulates Mre11 endonuclease activity. Proc Natl Acad Sci U S A 118, e2016287118.

Zhu, X. D., Küster, B., Mann, M., Petrini, J. H., and de Lange, T. (2000). Cell-cycle-regulated association of RAD50/MRE11/NBS1 with TRF2 and human telomeres. Nat Genet 25, 347–352.

Zinder, J. C., Olinares, P. D. B., Svetlov, V., Bush, M. W., Nudler, E., Chait, B. T., Walz, T., and de Lange, T. (2022). Shelterin is a dimeric complex with extensive structural heterogeneity. Proc Natl Acad Sci U S A 119, e2201662119.

